# Neuronal autophagosomes are transported to astrocytes for degradation

**DOI:** 10.1101/2024.09.03.610698

**Authors:** K. Linda, I.M.E. Schuurmans, H. Smeenk, K. Vints, M. Negwer, N. Peredo, E.I. Lewerissa, J. Swerts, M. Hoekstra, A. Mordelt, S. Kuenen, S.F. Gallego, F.M.S de Vrij, N. Nadif Kasri, P. Verstreken

## Abstract

Autophagy is a vital catabolic process responsible for the degradation of cytosolic components, playing a key role in cellular homeostasis and survival. At synapses, autophagy is crucial for regulating neuronal activity and utilizes a specialized machinery. While considerable progress has been made in understanding the initiation of autophagy and autophagosome formation, the mechanisms governing the clearance of autophagosomes from synaptic sites remain poorly understood. Here, we identify a novel pathway in which astrocytes actively participate in the clearance of pre-synaptic autophagosomes. Using neurons derived from human induced pluripotent stem cell (hiPSC) lines expressing fluorescent autophagy markers and chimeric mouse models, we demonstrate that neuronal autophagosomal vesicles are physically transferred to astrocytes, a process that is enhanced when synaptic activity is suppressed. Autophagosome transfer does not require direct physical cellular contact, but it does require Dynamin and cholesterol-dependent endocytosis for the internalized neuronal autophagosomes to ultimately fuse with astrocytic lysosomes. Our findings reveal a previously unrecognized mechanism of neuronal autophagosome clearance that does not require slow axonal retrograde transport but their transfer to nearby astrocytes.

## Introduction

Neurons, characterized by their complexity, polarization and compartmentalization, present with unique challenges to maintain proteostasis at axon endings due to the often-considerable distance between synapse and cell soma. Astrocytes play a crucial non-cell-autonomous role in supporting neurons by regulating the extracellular environment, providing essential metabolic resources, and clearing extracellular debris and neurotransmitters^1–4^. Neurons also employ cell-autonomous mechanisms to deal with the challenges posed by their polarized structure. These include axonal transport systems, localized signaling pathways, and the compartmentalization of specific organelles^5–7^, such as degradative autophagosomes, which collectively enable the efficient management of proteostasis, even at distal synaptic sites^8^.

Macroautophagy (referred to as autophagy) is a highly conserved evolutionary process that plays a crucial role in the catabolic degradation and clearance of cytosolic content, including long-lived proteins, protein aggregates, and redundant or damaged organelles^9^. During this process, autophagic cargo is encapsulated within cytosolic double-membraned organelles known as autophagosomes. These autophagosomes can subsequently fuse with lysosomes, facilitating the degradation and recycling of their contents. This process is vital for maintaining cellular homeostasis and cell viability under various conditions^10–12^ and is essential during cellular development and differentiation^13^.

Autophagy, in particular, appears to be regulated in a compartment-specific manner to ensure proper brain and neuron function^14,15,16^. Emerging evidence suggests the presence of a highly specialized machinery that governs autophagy at pre-synaptic sites, with regulation closely tied to neuronal activity and activity-controlled calcium flux ^17–21^. This indicates that synaptic autophagosomes are formed locally at synapses utilizing mechanisms that act independently from basal autophagy active in the soma. This allows pre-synaptic compartments to autonomously manage protein and organelle turnover. The current understanding is that these presynaptic autophagosomes then engage in a lengthy retrograde journey to the soma, during which they gradually acidify through lysosomal fusion, ultimately leading to the degradation and recycling of their contents^22^.

In this study, we demonstrate that neuronal autophagosomes can follow an alternative pathway by being transported to local astrocytes for degradation. Utilizing neurons derived from human induced pluripotent stem cell (hiPSC) lines and chimeric mice, we find that neuronally-produced autophagosomes are transported to and degraded by astrocytes. This process does not require physical cell-contact but does depend on clathrin-independent and dynamin- and cholesterol-dependent endocytosis. This local mechanism of neuronal autophagosome clearance provides an efficient alternative to the slower process of retrograde axonal transport and endows astrocytes with recycled biomolecules derived from degraded neuronal components.

## Results

### Neuronal LC3 accumulates in astrocytes

To investigate mechanisms of neuronal autophagy in human cells we used a control human induced pluripotent stem cell (hiPSC) line for which one allele of the *MAP1LC3B* gene was fused to *GFP*, enabling the endogenous expression of a GFP-tagged LC3, a protein that labels autophagosomes^23^. We differentiated these cells into cortical, excitatory neurons (iNeurons) through doxycycline-induced Neurogenin2-overexpression^24^ and co-cultured them for 4 weeks with non-tagged primary rat astrocytes (Figure 1A). While we observe clear LC3-labeled puncta (autophagosomes) in neurites, we were surprised to also find an abundance of GFP-LC3 puncta in the non-tagged astrocytes (Figure 1B).

**Figure 1:**
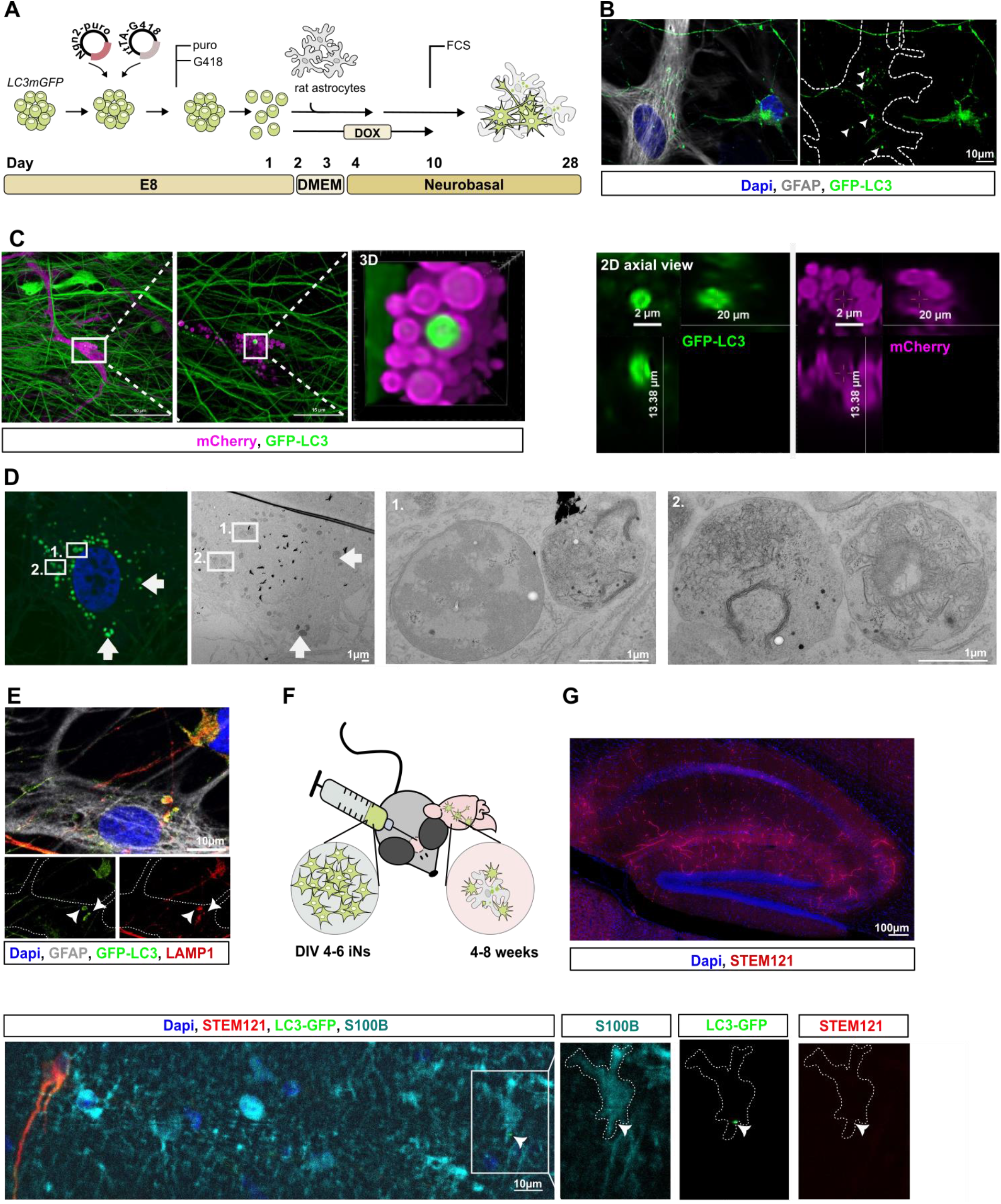
Neuronal LC3 accummulates in astrocytes. **A** Schematic representation of experimental setup. **B** Representative image of 4 week old iNeuron-astrocyte co-culture. **C** Life cell imaging including 3D and 2D axial view of GFP-LC3 puncta localized in mCherry positive astrocyte cultured with GFP-LC3 hiPSC-derived neurons after 4 weeks of differentiation. **D** CLEM images of 4 week of hiPSC-derived neuron including two representative regions. **E** Representative image showing GFP-LC3 (green) co-localizing with LAMP1 (red) in astrocytes marked with GFAP (grey). **F** Schematic representation of experimental setup and (**G**) representative image of STEM121 (red)-positive human neurons successfully transplanted into mouse brain. **H** Representative image of mouse brain slice showing a STEM121-positive (red) neuron on the left and GFP-LC3 puncta (green) within S100B-positive (cyan) astrocytic structures on the right.

We verified the unlabeled cells in which GFP-LC3 puncta appeared, were not unwanted “contaminating cells” derived from the hiPSC-differentiation and used a lentivirus to create mCherry-expressing rat astrocytes. Co-cultures of LC3-GFP-expressing iNeurons with mCherry-expressing astrocytes shows abundant GFP-LC3 puncta in red fluorescent cells, confirming the notion of transfer of the label from neurons to astrocytes (Figure 1C).

We ascertained these GFP-LC3 puncta in astrocytes are autophagosomes using correlative light and electron microscopy (CLEM). The GFP-positive puncta are spherical, vesicle-like structures with evident closed membranes. About 25% of the imaged vesicles present with a double membrane, reminiscent of classical autophagosomes while the remaining 75% of vesicles has a single membrane and include various electron dense structures, reminiscent of autolysosomes and other degradative organelles (Figure 1D). We confirm this by our observation that the LC3-GFP puncta in astrocytes co-localize with LAMP1, a protein that marks late endosomes and lysosomes (Figure 1E) and that many puncta also label with Lysotracker that stains acidic organelles (see below).

### In vivo accumulation of neuron-derived LC3 structures in astrocytes

To test if the transfer of neuronal LC3 to astrocytes also occurs *in vivo* we transplanted LC3-GFP-expressing early Ngn2-induced cells into *Rag2* ^*-/-*^ mouse brains (5 independent experiments; Fig 1F)^25^. We analyzed these samples between 4- and 8-weeks post-transplantation by labelling brain slices with Aldh1l1 and S100B (astrocyte markers), STEM121 (human cell marker) and anti-GFP (Figure 1H). This shows, as expected, the transplanted GFP-positive and STEM121-positive hiPSC-derived neurons, but also GFP-positive inclusions in Aldh1l1-/S100B-labeled astrocytes that are not positive for STEM121 (Figure 1G-H, Sup. Figure 1C). We find such GFP-labeled astrocytes consistently across 5 animals and they appear in regions close to the transplanted human neurons (Figure 1H). Taken together these data suggest LC3-GFP can be transferred from neurons to nearby astrocytes.

### Increased neuronal LC3 accumulation in astrocytes during neuronal silencing

We next asked which conditions alter the accumulation of neuron-derived LC3-structures in astrocytes (Figure 2A, Supplementary Table 1). We induced autophagy in our co-cultures using overnight treatment with Rapamycin (Rap) or Buthionine sulfoximine (BSO). While we showed these conditions activate neuronal autophagy^15^, the basal number of LC3-labeled puncta that appear in astrocytes does not change (Figure 2B). Hence, merely activating autophagy in neurons does not trigger additional LC3 transfer to astrocytes.

**Figure 2:**
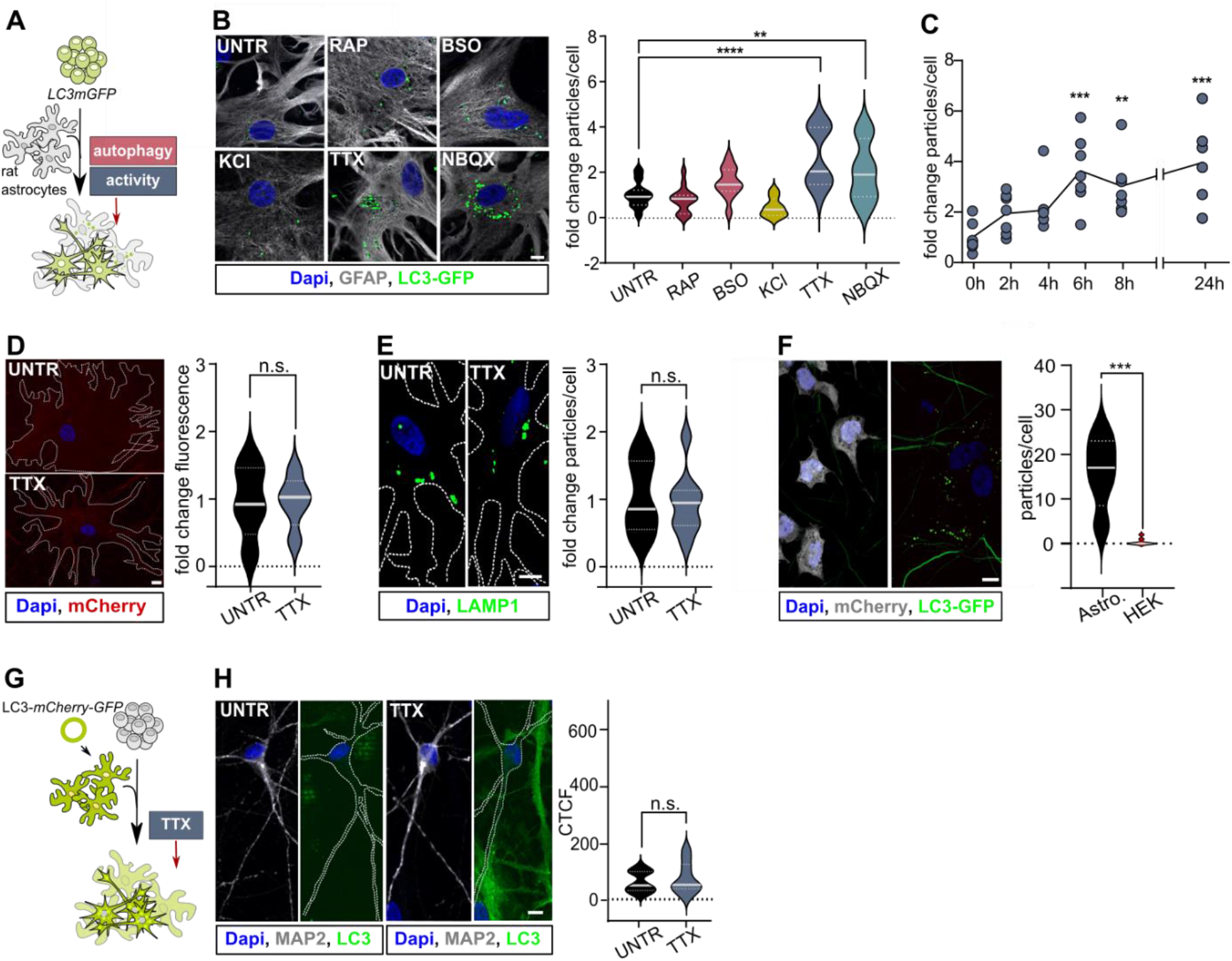
Increased neuronal LC3 accumulation in astrocytes during synaptic silencing. **A** Schematic presentation of the experimental setup. **B** Representative images of GFAP (grey) labelled astrocytes containing LC3-GFP (green) puncta, and puncta quantification after 4 weeks of co-culture with LC3-GFP iNeurons. Number of puncta was compared to the number of LC3 puncta in astrocytes of untreated (UNTR) cultures. N=3 **C** Quantification of LC3-GFP puncta in astrocytes co-cultured for four weeks with LC3-GFP iNeurons after 2, 4, 6, 8, and 24 hours TTX treatment. We compared puncta in TTX treated cultures to the number of puncta in astrocytes of untreated culture (0h). N=2 **D** Representative images of outlined astrocytes that were co-cultured four weeks with mCherry-positive iNeurons with and without overnight TTX treatment; mCherry signal in TTX treated astrocytes was compared to mCherry signal in untreated (UNTR) astrocytes. N=2 **E** Representative images of outlined astrocytes that were co-cultured four weeks with LAMP1-LC3 iNeurons with and without overnight TTX treatment; Number of GFP**-**Lamp1 puncta (green) in TTX-treated astrocytes was compared Lamp1-GFP puncta in untreated astrocytes. N=2 **F** Representative image of mCherry (grey)-positive HEK293T cells co-cultured with primary rat astrocytes and LC3-GFP iNeurons after TTX treatment. Particles were quantified in HEK cells and near astrocytic nuclei. **G** Schematic presentation of experimental setup **H** LC3-GFP-mCherry-infected astrocytes were co-cultured with control iNeurons and treated with TTX after differentiating for four weeks. GFP signal was quantified in iNeurons and compared to non-treated cells. N=2.

We then asked if neuronal activity modulates LC3-GFP transfer. We applied KCl that triggers membrane depolarization and increases neuronal activity, but this treatment also does not affect the number of puncta in astrocytes (Figure 2B). We subsequently blocked neuronal activity using NBQX, an AMPA receptor antagonist or using TTX, a Na^2+^ channel blocker that inhibits action potentials. Overnight treatment results in a significant increase in the number of LC3-GFP labeled particles per astrocyte (Figure 2B). We independently confirmed this result using another control hiPSC line that we infected with a lentivirus to express LC3-mCherry. Incubating co-cultures of this cell line with TTX also causes the increased number of LC3 puncta in astrocytes, indicating our initial finding is not cell line-dependent (Sup. Figure 1A). To further characterize our observation, we conducted a time-course experiment and find a significant increase of LC3-GFP puncta in astrocytes already after 6 h of TTX treatment. Furthermore, this effect remains stable up to 24 h after TTX was added (Figure 2C). Hence, silencing neuronal activity triggers the formation of neuron derived LC3-accumulations in astrocytes.

### Transfer of LC3 structures is specific for astrocytes

Our next question was to determine the specificity of the transferred LC3. We first asked if TTX triggered general protein transfer or if this was more specific for LC3 protein and/or structures. We used lentivirus to express untagged mCherry in hiPSCs and a hiPSC line for which one allele of the *LAMP1* gene was fused to GFP, differentiated the cells into neurons and incubated them with unlabeled astrocytes. While mCherry signal and LAMP1-GFP signal are detectable in astrocytes, overnight TTX treatment does not change these basal levels (Figure 2D-E). This is interesting because while most LC3 particles that appeared in astrocytes upon TTX treatment overlapped with LAMP1 staining (see Figure 1), this LAMP1 does not appear to be derived from neurons. Together, the results indicate that while there is some transfer of proteins between cells (mCherry, LAMP1), the accumulation of LC3 puncta in astrocytes upon TTX treatment is a specific mechanism.

To then test if the transfer of LC3 is specific from neurons towards astrocytes we first introduced mCherry-expressing HEK293T cells into our iNeuron-astrocyte co-cultures. Following TTX treatment we only found LC3-GFP puncta in astrocytes and almost none in the HEK 293T cells (Figure 2F). Second, we asked if LC3 could also be transferred from astrocytes to neurons upon silencing neuronal activity. We created (unlabeled) iNeuron and LC3-GFP-mCherry expressing astrocyte co-cultures and treated them with TTX (Figure 2G), but we do not find increased levels of mCherry-LC3 puncta in neurons (Figure 2H). These results indicate that there is directional transfer of neuronal LC3 to astrocytes (and potentially other cell types we did not (yet) test).

### LC3 transfer is mediated without cell-to-cell contact

To start investigating the mechanism of transfer of LC3 structures and/or protein we first asked if cell contact is required. Such mechanisms could involve close gap-like junctions or tunneling nanotubes^26–28^. We cultured LC3-GFP-expressing iNeurons and unlabeled astrocytes on two different coverslips and placed the coverslip with astrocytes upside down over the neurons, spaced by 1-2 mm thick Paraffin dots to avoid direct cell-cell contact^29^ (Figure 3A). TTX treatment again causes the significant increase in the number of LC3-GFP puncta in astrocytes (Figure 3B), a phenotype similar to the one we found with our integrated co-cultures. To further exclude the possibility that transfer was occurring via longer-range cell-cell connections that would bridge the gap created by the paraffin spacers, we proceeded by collecting medium from LC3-GFP expressing iNeuron co-cultures treated with TTX and then added this to unlabeled astrocyte monocultures. Also here, we observe the appearance of LC3-GFP particles in the astrocytes following a 3h incubation period (Figure 3C). These results demonstrate that direct cellular contact is not required for LC3 transfer to astrocytes.

**Figure 3:**
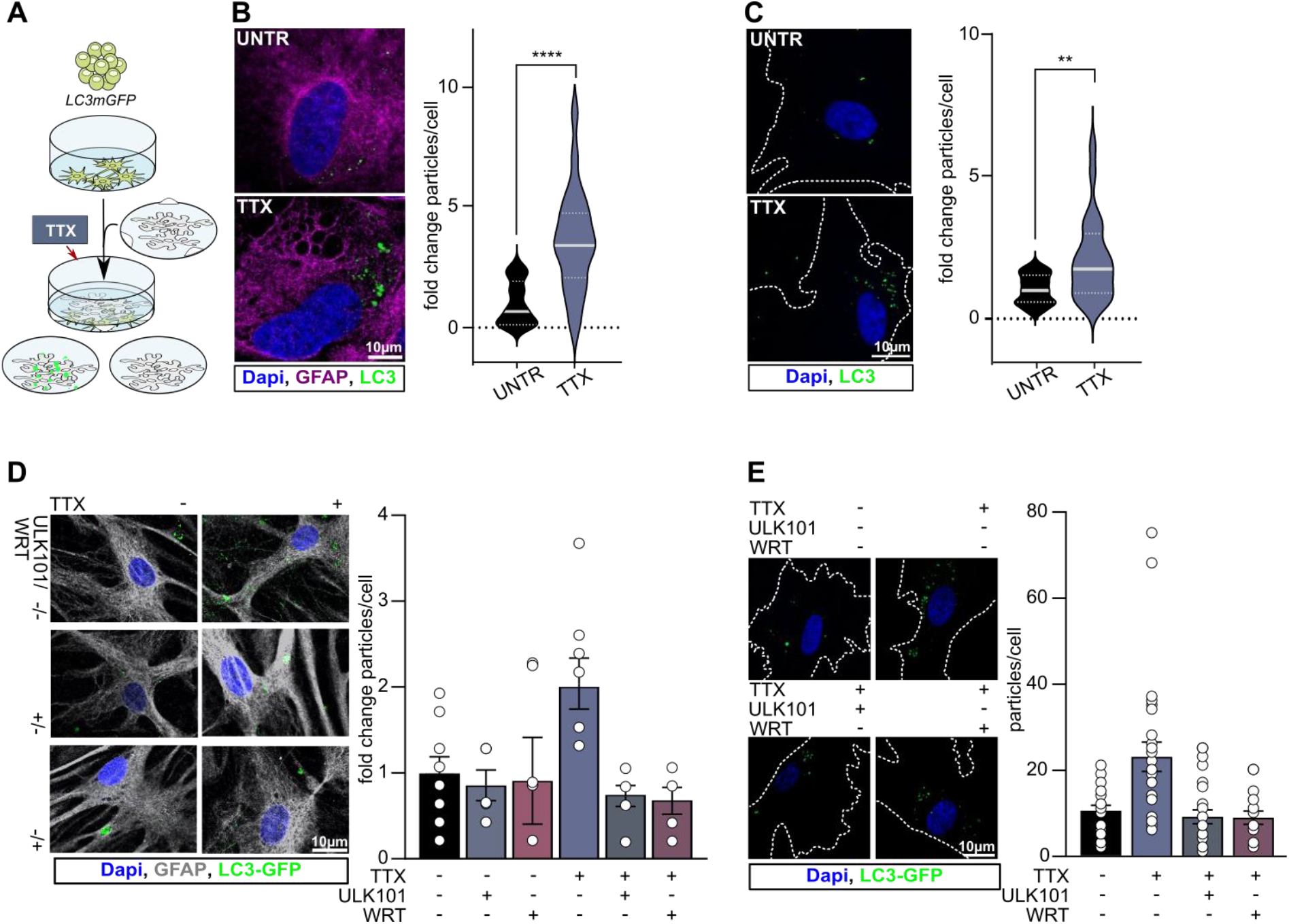
LC3 transfer mechanisms. **A** Schematic presentation of sandwich cultures, used to culture neurons and astrocytes in close proximity, but without cell-to-cell contact. **B** Representative images of sandwich cultured astrocytes with GFAP in magenta and LC3 in green. GFP-LC3 puncta were quantified in TTX treated astrocytes and compared to untreated (UNTR) cells. **C** Representative images of outlined astrocytes that were incubated with medium harvested after overnight TTX treatment of a LC3-GFP iNeuron-astrocyte co-culture. Quantification of the number of GFP particles/cell which was compared to the number of particles/cell in astrocytes that were incubated with medium of untreated cultures (UNTR). N=2 **D** Representative images of LC3-GFP iNeuron-astrocyte co-cultures after overnight TTX treatment in presence of autophagy blockers. LC3-GFP puncta (green) were quantified in Gfap-positive astrocytes (grey) and compared to untreated co-cultures. N=2 **E** Representative images of outlined astrocytes that were incubated with whole medium harvested after overnight TTX treatment of a LC3-GFP iNeuron-astrocyte co-culture in presence of autophagy blockers. GFP-LC3 particles were quantified per cell. N=3.

To characterize the transferred LC3-GFP moieties we used Amplicon ultracentrifugation columns to isolate a >10kDa and a >100kDa medium fraction (Sup. Figure 2). Unlipidated LC3-GFP is 44kDa and monomers and are expected to reside in the >10kDa fraction and to be excluded from the >100kDa fraction. First, we detect GFP signal in both fractions; second, using negative stain EM we find vesicular structures in both fractions but the >10 kDa fraction does not include larger vesicles within the size range of an autophagosome, which we find in the >100 kDa fraction (Sup. Figure 2). We incubated astrocytes with these fractions and find LC3-GFP accumulations in either condition, but the >100kDa fraction is more potent than the >10kDa fraction (Sup Figure 2B). These results suggest that larger LC3-decorated membrane-structures can be transferred to astrocytes.

We challenged our notion that autophagosomal vesicles are being transferred in their entirety using a genetics experiment involving astrocytes in which the core-autophagy component Atg5 was knocked down^30^. We confirm that these astrocytes (in monocultures) cannot form (endogenous) autophagosomes (Sup. Figure 2C). However, when we add medium from LC3-GFP expressing iNeuron co-cultures treated with TTX, we observe the appearance of LC3-GFP-labeled structures inside these autophagy-defective astrocytes (Sup. Figure 2D). These data further support our conclusion that entire autophagosomal vesicles are being transferred from neurons to astrocytes.

### Autophagy in neurons is required for activity-dependent LC3 transfer to astrocytes

We reasoned that if entire autophagosomal vesicles are transferred (and not merely LC3 protein) that blocking neuronal autophagosome biogenesis during overnight TTX treatment should trump the appearance of LC3-GFP structures in astrocytes. We created LC3-GFP-expressing iNeuron astrocyte co-cultures and during TTX treatment incubated them with ULK-101, an inhibitor of autophagy induction or Wortmannin (WRT), that inhibits autophagosome formation. While these treatments do not affect overall LC3-GFP protein expression, either compound blocks the increased LC3 transfer to astrocytes (Figure 3D). We independently confirm this by also harvesting medium of these co-cultures and adding this to unlabeled astrocytes. Medium from TTX and ULK-101 or WRT-treated cultures failed to show the increased number of LC3-GFP labeled structures in astrocytes that we see with medium from TTX-only treated cultures (Figure 3E). These data are in further support of a model where vesicular structures are transferred and indicate that the increased transfer of LC3-positive vesicles following TTX stimulation relies on active autophagy in neurons.

### Astrocytes mediate LC3 vesicle uptake via clathrin-independent endocytosis

We wondered how astrocytes could incorporate neuronal LC3 decorated vesicles and assessed a role for an endocytic mechanism. Labeling astrocytes in TTX-treated sandwich-cultures reveals that the LC3-GFP-labeled structures co-localize with EEA1, a marker for early endosomes, and RAB7, a marker for late endosomes (Figure 4A), consistent with them moving through the endocytic trafficking pathway. To examine subsequent fusion with lysosomes, we first incubated astrocytes with Lysotracker-Red that stains acidic organelles like lysosomes, removed the dye and incubated the labeled cells with medium from TTX-treated co-cultures for 3h. Representative images shown in Figure 4A show that almost all LC3-GFP puncta co-localize with lysotracker, suggesting that endocytosed LC3-positive vesicles fuse with astrocytic lysosomes.

**Figure 4:**
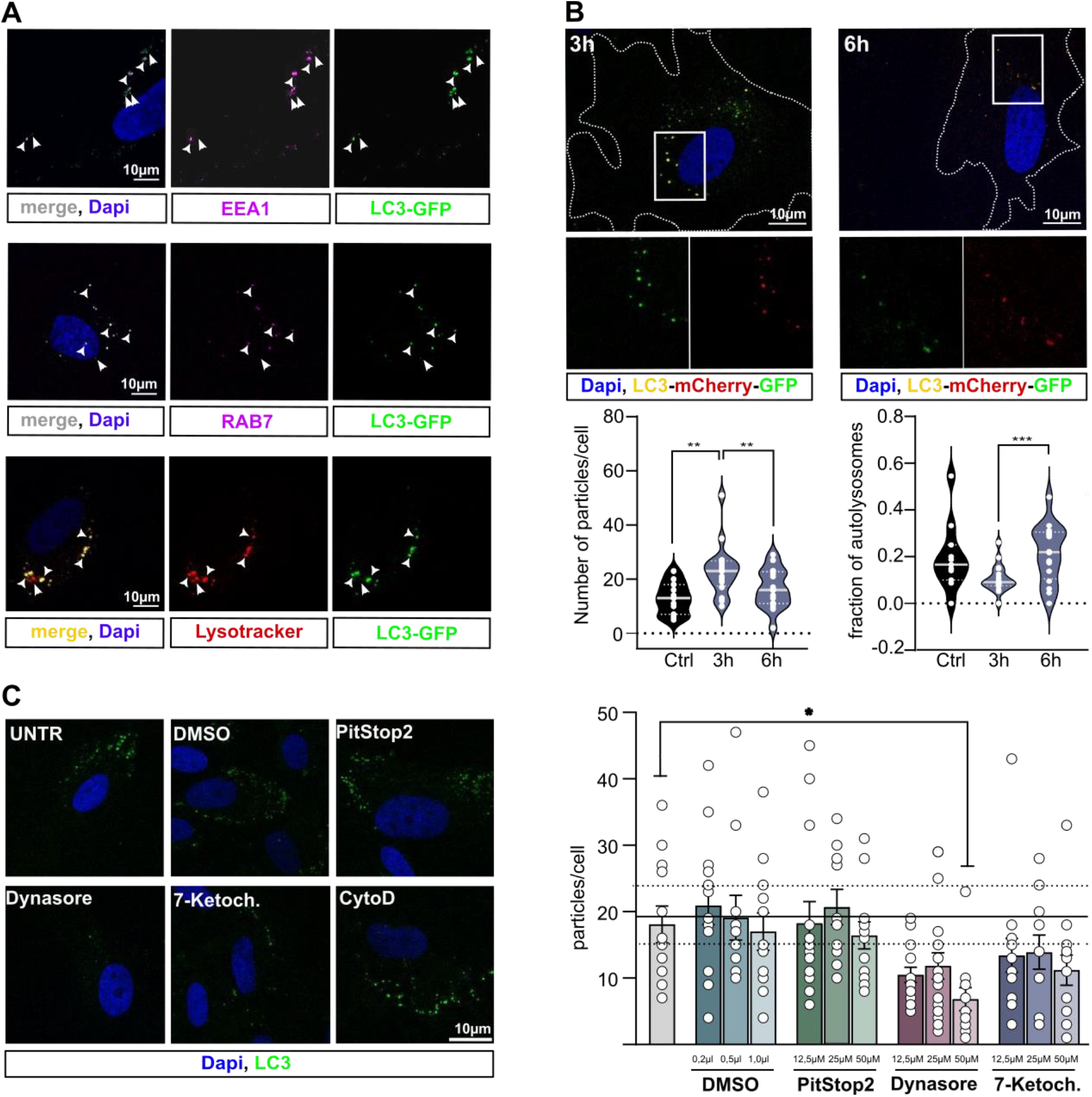
Astrocytes mediate autophagosomal uptake via clathrin-independent endocytosis. **A** Representative images of astrocytes of overnight TTX treated sandwich cultures with LC3-GFP iNeurons. Astrocytes were stained for EEA1 (magenta), RAB7 (magenta). Additionally, astrocyte monocultures were pre-incubated with lysotracker-red before the addition of medium harvested from TTX treated LC3-GFP iNeuron-astrocyte co-cultures. **B** Representative images of astrocytes that were incubated with medium harvested after overnight TTX treatment of a LC3-mCherry-GFP iNeuron-astrocyte co-culture. The number of neuronal LC3 puncta (red and yellow) was quantified after 3h and after 6h incubation and compared to astrocytes incubated for 3h with medium of an untreated co-culture (Ctrl). In the same images the fraction of autolysosomes (red) was quantified and compared to the Ctrl condition. N=2 **C** Representative images of astrocytes that were incubated with medium harvested after overnight TTX treatment of a LC3-GFP iNeuron-astrocyte co-culture in the presence of different endocytosis blockers. LC3-particles (green) per cell were quantified and compared to untreated astrocytes that were incubated with the same medium. N=2.

We then conducted time-course experiments, reasoning that the LC3-GFP vesicles that are internalized into astrocytes ultimately fuse with lysosomes for degradation (also predicted by our CLEM results (see Figure 1)). To this end we made use of the LC3-GFP-mCherry lentiviral construct, which is a widely used tool to monitor autophagic degradation. The GFP signal is pH-sensitive and degrades in the acidic environment of the autolysosome, while the mCherry signal is stable. This dual fluorescence enables the distinction between autophagosomes, show both green and red signals, and acidic autolysosomes, only showing red fluorescence^31^. We differentiated LC3-GFP-mCherry transduced iNeurons in co-culture with astrocytes for four weeks and then treated those cultures overnight with TTX. The harvested medium was then added to astrocytes for 3 hours and 6 hours. Quantification of fluorescent signal after 3 hours showed an elevated number of neuronal LC3 puncta (green and red) when compared to astrocytes incubated with medium of untreated cultures (Figure 4B). Interestingly, that number reduces after 6 hours. At the same time we observe an increase in the fraction of red puncta at this time point, suggestive for the formation of autolysosomes. This is consistent with the internalized LC3-GFP-labeled structures to enter a degradative endocytic trafficking pathway in astrocytes.

To reveal the mechanism of LC3-GFP-vesicle endocytosis in astrocytes we used different endocytosis blockers (Supplementary Table 2). Astrocytes pre-treated with these blockers were incubated for 3 hours with medium from TTX treated co-cultures. Cytochalasin D induces toxicity precluding us from drawing conclusions (Supplementary Figure 3A-B), while PitSTOP2, even at the highest concentrations, does not affect the uptake of neuronal LC3 (Figure 4C), suggesting Clathrin-mediated endocytosis is not involved.

We then tested Dynasore, a blocker of dynamin-dependent endocytosis, and 7-Ketocholesterol that inhibits cholesterol-dependent endocytosis such as caveolar invaginations^32^. Preincubation of astrocytes with increasing concentrations of either compound inhibits the increased appearance of LC3-labeled structures in astrocytes. Although endocytosis blocker lack specificity and exhibit various side effects^33–35^, the results provide preliminary evidence that internalization of neuron-derived LC3-decorated vesicles into astrocytes requires a Clathrin-independent and Dynamin- and cholesterol-dependent endocytic process, providing the starting point for more targeted genetic inhibition approaches in the future.

## Discussion

We demonstrate that in human neurons, LC3-labeled autophagosomal vesicles can be selectively transferred to astrocytes for degradation. This transfer process is negatively regulated by neuronal activity and requires Clathrin-independent, Dynamin-, and/or cholesterol-dependent endocytosis in astrocytes for the autophagosomes to be delivered to astrocytic lysosomes. This mechanism is particularly significant in the context of synapses, where lysosomal fusion and autophagosome acidification primarily occur during the lengthy transport to the soma, with less acidification at the synapses themselves^36,37^. Our findings suggest that the close proximity of astrocytes offers an alternative route: local transport to and recycling within astrocytes. This process is especially relevant for long-lived, non-dividing neurons and may be critical in neurodegenerative conditions, where synapses are vulnerable, and protein aggregates tend to accumulate, disrupting local proteostasis^38^.

Our findings align with the concept of secretory autophagy, where double-membraned autophagosomes fuse with the plasma membrane to secrete their contents, contributing to proteostasis^39^. This mechanism offers an efficient way for synapses to expel debris rapidly. We observed an increased transfer of autophagosomal vesicles specifically during synaptic silencing (TTX, NBQX), suggesting that synaptically formed autophagosomes are likely degraded through this process, although autophagosomes from other neuronal regions may also utilize this pathway. Future isolation and analysis of GFP-labeled vesicles during transport could provide further insights into their (synaptic) origin. Furthermore, additional analyses will be needed to understand the regulatory mechanisms that govern the increased transfer of autophagosomal vesicles specifically during synaptic silencing.

Secretory autophagy has been linked to synaptic remodeling^40^ and the secretion of aggregation-prone proteins, such as intracellular amyloid-beta and huntingtin, which play roles in neurodegenerative diseases^41^. While significant progress has been made in understanding the cargo and cellular mechanisms involved in secretory autophagy, the fate of these secreted autophagosomal vesicles remains largely unexplored. In this study, we show that astrocytes are key recipients of these vesicles, and the roles of other cell types such as microglia and oligodendrocytes remain to be investigated. However, in our *in vivo* xenotransplantation experiments, we observed most of the secreted LC3-labeled particles to be within or very close to astrocyte markers, suggesting the preferential targeting of these cells. Given the known roles of astrocytes, we propose that, beyond breaking down and recycling neuronal debris, this process may also function as a signaling pathway, conveying information about neuronal status and health.

Reducing synaptic activity triggers homeostatic plasticity in neurons, leading to adjustments in synaptic strength to maintain network stability^42^. This involves dynamic structural reorganization of synapses, with autophagy and astrocytes playing key roles^43–45^. While active autophagy facilitates remodeling of synaptic structures by removing and recycling obsolete or damaged synaptic proteins, astrocytes support the process by regulating extracellular ion balance, releasing neurotrophic factors, and modulating neurotransmitter levels^44–46^. Our work establishes a direct link between astrocytes and the processing of neuronal autophagosomal vesicles. We hypothesize that this transfer is crucial for the resilience and adaptability of neural circuits, but this warrants further investigation.

We found that astrocytes internalize LC3-labeled vesicles through a Clathrin-independent, but Dynamin- and cholesterol-dependent endocytic mechanism. Macropinocytosis, a Clathrin-independent pathway, mediates the nonspecific uptake of extracellular fluids, macromolecules, and cell fragments^47^. This process involves the formation of cup-like structures that extend from the cell surface, driven by the actin cytoskeleton, and that close around extracellular material to form micron-sized vesicles. These vesicles are then processed and recycled through the endocytic system^47^. Primary mouse astrocytes were shown to be able to take up extracellular vesicles via micropinocytosis^48^. However, many other cell types, including HEK293 cells that we tested here, also exhibit macropinocytosis. Interestingly, our observation that HEK293 cells failed to take up neuron-derived LC3-labeled autophagosomal vesicles after TTX stimulation suggests that autophagic vesicles are unlikely to be internalized through macropinocytosis. In future research it will be interesting to identify specific molecular targeting mechanisms that drive autophagic vesicle-recognition by astrocytes (and possibly other cell types).

Caveolar endocytosis is also a Clathrin-independent, Dynamin- and cholesterol-dependent process and is initiated by ligand binding, thus enabling targeted and controlled endocytosis^49^. This pathway involves omega-shaped plasma membrane invaginations (caveolae) and Dynamin-dependent internalization of cargo. Internalized caveolae can follow the classical endocytic route depending on the ligand^50^. Caveolae-mediated endocytosis showed to play a role in TLR4 endocytosis in cortical astrocytes^51^. However, while our findings show extensive co-localization of LC3-GFP puncta with late endosomal and lysosomal markers (Figure 1E, 4A), endocytosed TLR4 exhibits a slower and less comprehensive transition to lysosomes. Another astrocytic endocytosis pathway, which aligns more closely with this aspect, is Clathrin- and Dynamin-independent and regulated by Rab5^52^. This pathway is characterized by the rapid transition of endocytosed vesicles, from early to late endosomes and lysosomes. Such a pathway then likely facilitates the fast degradation and recycling of neuronal autophagic vesicles, which may be essential for astrocytes to respond rapidly to changes in synaptic activity and maintain neuronal circuit adaptability.

## Materials & Methods

### hiPSC lines and culture

We used two independent male control hiPSC lines. Both cell lines originate from reprogrammed fibroblasts. The first line was obtained from Coriell (AICS-0030-022:WTC-mEGFP-MAP1LC3B-cl22). For this line one allele of the *MAP1LC3B* gene was endogenously tagged with *mGFP*, so that fluorescently labelled LC3 protein was generated. The second control line was obtained from the Tau Consortium Stem Cell Group^53^. Additionally we used hiPSC lines of the same genetic background as the *MAP1LC3B*-*GFP* hiPSC line, either unlabelled (GM25256) or with an endogenously GFP tagged *LAMP1* gene (AICS-0022-037: MONO-ALLELIC mEGFP-TAGGED LAMP1 WTC iPSC LINE (TAG AT C-TERM)). All clones used in this study were validated through a battery of quality control tests including-morphological assessment and karyotyping to confirm genetic integrity. All clones expressed the stem cell markers POU5F1, SOX2 NANOG, SSEA-4, and TRA-1-81.

Cells were cultured on Geltrex coated wells in E8 Flex medium supplemented with Pen/Strep (Gibco). Medium was refreshed every other day and cells were split twice per week.

### Astrocyte culture

Primary rat astrocytes were isolated from rat brain as described previously^24^. In short cortices were dissected from rat pubs (E18 or P0-2). After dissociating the cells were cultured at 37°C, 5%CO2 in DMEM high glucose (Gibco) supplemented with 10% FBS (Gibco) and 1% Pen/Strep for at least one week with regular medium changes and shaking to remove less adherent cells (i.e. microglia). When not used freshly, astrocytes were dissociated from T75 flasks with trypsin and resuspended in cryopreservation medium and stored in liquid nitrogen after two passages. After thawing the astrocytes were passaged or collected for experimental use when reaching 80% confluency. When plated for transfer experiments astrocytes were plated on PLO/human Laminin coated coverslips in DMEM high glucose supplemented with 10% FBS and 1% Pen/Strep. Two to three days after plating medium was switched to neurobasal medium supplemented with B27 (Gibco), Glutamax (Gibco), 1% Pen/Strep, NT3 (Promokine), BDNF (Promokine) and 2.5% FBS. All experiments were performed with at least two independent batches of primary rat astrocytes.

### Neuronal differentiation

iPSCs were differentiated towards cortical excitatory neurons as described previously^24^. Summarized, lentiviral vectors were used to stably integrate transgenes encoding for rtTA and doxycycline inducible Ngn2. To select for iPSCs that were transduced with both lentiviral vectors we started G418 (0.5 μg/ml; Sigma-Aldrich) and puromycin (0.1 ug/ml; Sigma-Aldrich) selection 48 h after infection. The antibiotics concentration was doubled at day two and three of the selection process. iPSCs surviving the selection process were cultured at general iPSC culture conditions, E8 Flex medium was additionally supplemented with low concentration puromycin and G418.

To start the differentiation process stabilized iPSC lines were singularized using Accutase and plated at a density of 100 cells/mm^2^ in E8 basal flex medium (Gibco) supplemented with Pen/Strep, RevitaCell, and doxycycline (1 μg/ml) on poly-L-ornithine (50 μg/mL; Sigma-Aldrich) and human laminin (Biolamina LN521) coated wells with or without coverslips.

The day after plating medium was changed to DMEM-F12 (Gibco) supplemented with N2 (Gibco), NT3 (Promokine), BDNF (Promokine), MEM non-essential amino acid solution NEAA (Sigma-Aldrich, M7145), doxycycline (2 μg/ml), and Pen/Strep. To support neuronal maturation primary rat astrocytes were added to the culture in a 1:1 ratio two days after plating. At DIV 3 the medium was changed to Neurobasal medium (Gibco) supplemented with B-27, Glutamax, Pen/Strep, NT3, BDNF, and doxycycline (1 μg/ml). Cytosine β-D-arabinofuranoside (Ara-C) (2 μM; Sigma-Aldrich) was added once to remove proliferating cells from the culture. From DIV 6 onwards half of the medium was refreshed three times a week. Addition of 1 μg/ml doxycycline was stopped after two weeks. The medium was additionally supplemented with 2.5% heat inactivated FBS to support astrocyte viability from DIV10 onwards. Neuronal cultures were kept through the whole differentiation process at 37°C, 5%CO2.

### Immunocytochemistry

After 4 weeks of differentiation and respective treatments, cells were fixed with 4% paraformaldehyde, 4% sucrose (v:v) and permeabilized with 0.2% Triton X-100 (Sigma-Aldrich, T8787) in PBS (Sigma-Aldrich, P5493) for 10 min. Aspecific binding sites were blocked by incubation in blocking buffer (PBS, 5% normal goat serum (Invitrogen), 1% bovine serum albumin (Sigma-Aldrich, A7906), 1% glycine (Sigma-Aldrich, G5417), 0.2% Triton X-100 (Thermo Fisher)) for 1 h at room temperature (RT). Primary antibodies (GFAP, Aveslabs #GFAP 1:500; Synapsin1, SanBio 51-5200 1:1000; Lamp1, Sigma-Aldrich L1418 1:200; EEA1, Fisher Scientific MA5-14794 1:200; Rab7, Sigma Aldrich R8779 1:1000; Map2,Sanbio 188004 1:1000) were diluted in blocking buffer for overnight incubation at 4°C. Secondary antibodies conjugated to Alexa Fluor-fluorochromes (goat-anti rabbit Alexa647, Invitrogen A21245; goat-anti rabbit Alexa555, Invitrogen A21429; goat-anti mouse Alexa555, Invitrogen A21424; goat-anti mouse Alexa647, Invitrogen A21236; goat-anti chicken Alexa647, Invitrogen A21472), were diluted 1:2000 in blocking buffer and applied for 1 h at RT. Hoechst was used to stain the nucleus before cells were mounted using DAKO or Mowiol fluorescent mounting medium and stored at 4°C. Samples were imaged with Nikon NiE A1R confocal miscroscope.

### Correlative Light and Electron Microscopy (CLEM)

22000 iPSCs were plated on PLO/ human laminin coated 3cm Mattek glass bottom dishes and differentiated according to above described differentiation protocol. After 4 weeks differentiation was stopped by adding double concentrated fixative 8% PFA + 0,4% GA in 0,1M PB (pH 7.4) to the medium (1:1), after 10 minutes the solution was replaced by single concentrated fixative 4% PFA + 0,2% GA in 0,1M PB (pH 7.4) for 2 hours incubation at room temperature in the dark. After this pre-fixation the sample was rinsed with 0.1M PB and imaged with Zeiss LSM980 Airyscan. Imaged regions were marked for correlation. Immediately after imaging, samples were fixed in 2,5% GA in 0,1M sodium cacodylate trihydrate buffer (pH 7,6) and prepared for electron microscopy.

After overnight incubation at 4 degrees samples were rinsed with 0,1M sodium cacodylate trihydrate buffer (pH 7,6), 3 × 7 min, on ice, and then incubated with 1% osmium tetroxide, 1.5% potassium ferrocyanide (K4Fe(CN)6) in 0.1M sodium cacodylate trihydrate buffer (pH 7.6) for 60 minutes on ice in the dark. This was followed by an overnight incubation at 4 degrees in the dark with 0.5% uranylacetate in 25 % methanol. Next samples were incubated en bloc with lead aspartate (Walton’s lead aspartate: 0.02 M lead nitrate in 0.03 M sodium aspartate, pH 5.5) for 30 minutes at 60 degrees. A graded series of ethanol: 1 × 10 min each step (30, 50, 70, 80, 95% ethanol) followed by two times 10 minutes pure ethanol, all on ice were applied for dehydration. This was followed by infiltration and embedding with resin. Samples were polymerized for 48 hours at 60 degrees by putting BEEM® Embedding Capsules Size 3 upside down on the marked location of the cells of interest. The capsule was removed from the glass bottom by switching it between liquid nitrogen and 80° heating plate. The resin block was trimmed around the cells of interest which were recognized by their morphology. 70nm consecutive sections were cut with a Leica Ultracut S and collected on slot grids. The cells were imaged with a TEM (JEM1400-LaB6, JEOL) operated at 80kV, equipped with an Olympus SIS Quemesa 11 MP camera, at 1kx and 10kx magnification.

### Xenotransplantation experiments

All animal experiments were approved by the local animal welfare committee. Xenotransplantation was performed as described before^54^. NGN2+/rtTA+ iPSCs of the WTC-mEGFP-MAP1LC3B-cl22 line were differentiated to neurons by doxycycline induction (250 μg/ml) for 4 days, without addition of astrocytes. These neurons were then transplanted in immunodeficient neonatal Rag2-/-BALB/c x C57Bl/6NCrl F1 hybrid mice at P1 under cryoanesthesia. During the procedure, five 1μl injections were performed using a pulled glass pipette, bilaterally in the anterior and posterior anlagen of the corpus callosum, and one in the cerebellar peduncle. Approximately 5-10 × 10^4^ neurons were injected per injection site, in PBS with Fast Green FCF (Sigma-Aldrich, #F7252). Mice were sacrificed between 4 and 8 weeks after xenotransplantation, through transcardiac perfusion with saline followed by 4% paraformaldehyde (PFA) in PBS.

Brains were harvested and post-fixed for two hours in 4% PFA, after which they were incubated in 10% sucrose overnight at 4°C. The following day, brains were embedded in 12% gelatin/10% sucrose solution and fixed for another 2 hours in a 10% PFA/30% sucrose solution. They were then stored overnight in 30% sucrose solution at 4°C, after which they were sliced into 40 μm slices on a freezing microtome (Leica, #SM2000R).

### Immunohistochemistry

For immunohistochemical analyses, mouse brain slices were pre-incubated with blocking buffer (10% normal horse serum (ThermoFisher, #16050122), 0.5% Triton X-100 (Sigma-Aldrich, #T8787) in PBS). Afterwards, primary antibody staining (ALDH1L1, Abcam #ab871170 1:250; S100B, Synaptic Systems #287006 1:250; GFP, Abcam #ab13970 1:1000; Stem121, Takara #Y40410 1:500) was performed in staining buffer (2% normal horse serum, 0.5% Triton X-100 in PBS) for 48 hours at 4°C. Secondary antibody staining (CY3-AffiniPure Donkey anti-Mouse IgG, #715165151; Alexa Fluor 488-AffiniPure Donkey anti-Chicken IgG, #703545155; Alexa Fluor 647-AffiniPure Donkey Anti-Rabbit IgG #711605152; all Jackson Immunoresearch) was also performed in staining buffer for 2 hours at room temperature, followed by mounting in Mowiol 4-88 (Merck, 81381). Samples were imaged on a Zeiss LSM 800 confocal microscope.

### Particle quantification & Statistics

FIJI software was used to quantify particles in astrocytes. Threshold to detect fluorescent particles was set for each experiment individually for an image of untreated cells and remained the same throughout the different conditions tested within this specific experiment. GFAP-positive cells containing at least one fluorescent particle were selected. To reduce the chance that background signal is quantified particles were only count when they were at least 0.2 μm^2^ or bigger. Statistical analysis of the obtained data was performed using GraphPad Prism 8 (GraphPad Software, Inc., CA, USA). We first determined whether data were normally distributed. We tested statistical significance for different experimental conditions by one-way ANOVA. Individual samples were compared using Sidak’s or Dunnett’s multiple comparisons test. When only two conditions were compared, we used unpaired t test. For not normally distributed data, we applied Kruskal-Wallis test combined with Dunn’s or Sidak’s multiple comparison test. Results with P values lower than 0.05 were considered as significantly different (*), P < 0.01 (**), P < 0.001 (***), P < 0.0001 (****).

## Acknowledgements

This work was supported by EMBO (postdoctoral fellowship EMBO ALTF 509-2022 awarded to K.L.) and the ZonMW PSIDER program TAILORED (grant 10250022110002 to N.N.K. and F.M.S.V.).

## References

1. Eroglu, C. & Barres, B. A. Regulation of synaptic connectivity by glia. Nature 468, 223–231 (2010).

2. Herrera Moro Chao, D. et al. Hypothalamic astrocytes control systemic glucose metabolism and energy balance. Cell Metab 34, 1532-1547.e6 (2022).

3. Morizawa, Y. M. et al. Reactive astrocytes function as phagocytes after brain ischemia via ABCA1-mediated pathway. Nat Commun 8, 28 (2017).

4. Mahmoud, S., Gharagozloo, M., Simard, C. & Gris, D. Astrocytes Maintain Glutamate Homeostasis in the CNS by Controlling the Balance between Glutamate Uptake and Release. Cells 8, 184 (2019).

5. Guedes-Dias, P. & Holzbaur, E. L. F. Axonal transport: Driving synaptic function. Science (1979) 366, (2019).

6. Donato, A., Kagias, K., Zhang, Y. & Hilliard, M. A. Neuronal sub-compartmentalization: a strategy to optimize neuronal function. Biological Reviews 94, 1023–1037 (2019).

7. Giandomenico, S. L., Alvarez-Castelao, B. & Schuman, E. M. Proteostatic regulation in neuronal compartments. Trends Neurosci 45, 41–52 (2022).

8. Decet, M. & Verstreken, P. Presynaptic Autophagy and the Connection With Neurotransmission. Front Cell Dev Biol 9, (2021).

9. Klionsky, D. J. & Codogno, P. The Mechanism and Physiological Function of Macroautophagy. J Innate Immun 5, 427–433 (2013).

10. Yorimitsu, T. & Klionsky, D. J. Autophagy: molecular machinery for self-eating. Cell Death Differ 12, 1542–1552 (2005).

11. Klionsky, D. J. & Emr, S. D. Autophagy as a Regulated Pathway of Cellular Degradation. Science (1979) 290, 1717–1721 (2000).

12. Klionsky, D. J. The molecular machinery of autophagy: unanswered questions. J Cell Sci 118, 7–18 (2005).

13. Mizushima, N. & Levine, B. Autophagy in mammalian development and differentiation. Nat Cell Biol 12, 823–830 (2010).

14. Uytterhoeven, V. et al. Hsc70-4 Deforms Membranes to Promote Synaptic Protein Turnover by Endosomal Microautophagy. Neuron 88, 735–748 (2015).

15. Linda, K. et al. Imbalanced autophagy causes synaptic deficits in a human model for neurodevelopmental disorders. Autophagy 18, 423–442 (2022).

16. Maday, S. & Holzbaur, E. L. F. Compartment-Specific Regulation of Autophagy in Primary Neurons. J Neurosci 36, 5933–5945 (2016).

17. Soukup, S. F. et al. A LRRK2-Dependent EndophilinA Phosphoswitch Is Critical for Macroautophagy at Presynaptic Terminals. Neuron 92, 829–844 (2016).

18. Vanhauwaert, R. et al. The SAC1 domain in synaptojanin is required for autophagosome maturation at presynaptic terminals. EMBO J 36, 1392–1411 (2017).

19. Reimer, R. J. et al. Bassoon Controls Presynaptic Autophagy through Atg5. Neuron (2017) doi:10.1016/j.neuron.2017.01.026.

20. Okerlund, N. D. et al. Bassoon Controls Presynaptic Autophagy through Atg5. Neuron 97, 727 (2018).

21. Vijayan, V. & Verstreken, P. Autophagy in the presynaptic compartment in health and disease. Journal of Cell Biology Preprint at 10.1083/jcb.201611113 (2017).

22. Cason, S. E. & Holzbaur, E. L. F. Axonal transport of autophagosomes is regulated by dynein activators JIP3/JIP4 and ARF/RAB GTPases. Journal of Cell Biology 222, (2023).

23. Kabeya, Y. LC3, a mammalian homologue of yeast Apg8p, is localized in autophagosome membranes after processing. EMBO J 19, 5720–5728 (2000).

24. Frega, M. et al. Rapid Neuronal Differentiation of Induced Pluripotent Stem Cells for Measuring Network Activity on Micro-electrode Arrays. Journal of Visualized Experiments (2017) doi:10.3791/54900.

25. Linaro, D. et al. Xenotransplanted Human Cortical Neurons Reveal Species-Specific Development and Functional Integration into Mouse Visual Circuits. Neuron 104, 972-986.e6 (2019).

26. Fróes, M. M. et al. Gap-junctional coupling between neurons and astrocytes in primary central nervous system cultures. Proceedings of the National Academy of Sciences 96, 7541–7546 (1999).

27. Sun, X. et al. Tunneling-nanotube direction determination in neurons and astrocytes. Cell Death Dis 3, e438–e438 (2012).

28. Zurzolo, C. Tunneling nanotubes: Reshaping connectivity. Curr Opin Cell Biol 71, 139–147 (2021).

29. Ioannou, M. S., Liu, Z. & Lippincott-Schwartz, J. A Neuron-Glia Co-culture System for Studying Intercellular Lipid Transport. Curr Protoc Cell Biol 84, (2019).

30. Mizushima, N. et al. A protein conjugation system essential for autophagy. Nature 395, 395– 398 (1998).

31. Leeman, D. S. et al. Lysosome activation clears aggregates and enhances quiescent neural stem cell activation during aging. Science (1979) 359, 1277–1283 (2018).

32. Lyons, M. A. & Brown, A. J. 7-Ketocholesterol. Int J Biochem Cell Biol 31, 369–375 (1999).

33. Willox, A. K., Sahraoui, Y. M. E. & Royle, S. J. Non-specificity of Pitstop 2 in clathrin-mediated endocytosis. Biol Open 3, 326–331 (2014).

34. Preta, G., Cronin, J. G. & Sheldon, I. M. Dynasore - not just a dynamin inhibitor. Cell Communication and Signaling 13, 24 (2015).

35. Szewczyk-Roszczenko, O. K. et al. The Chemical Inhibitors of Endocytosis: From Mechanisms to Potential Clinical Applications. Cells 12, 2312 (2023).

36. Maday, S., Wallace, K. E. & Holzbaur, E. L. F. Autophagosomes initiate distally and mature during transport toward the cell soma in primary neurons. Journal of Cell Biology 196, 407– 417 (2012).

37. Maday, S. & Holzbaur, E. L. F. Autophagosome Biogenesis in Primary Neurons Follows an Ordered and Spatially Regulated Pathway. Dev Cell 30, 71–85 (2014).

38. Tseng, C.-S., Chao, Y.-W., Liu, Y.-H.Huang, Y.-S. & Chao, H.-W. Dysregulated proteostasis network in neuronal diseases. Front Cell Dev Biol 11, (2023).

39. Ponpuak, M. et al. Secretory autophagy. Curr Opin Cell Biol 35, 106–116 (2015).

40. Chang, Y.-C. et al. Identification of secretory autophagy as a mechanism modulating activity-induced synaptic remodeling. Proceedings of the National Academy of Sciences 121, (2024).

41. Gonzalez, C. D., Resnik, R. & Vaccaro, M. I. Secretory Autophagy and Its Relevance in Metabolic and Degenerative Disease. Front Endocrinol (Lausanne) 11, (2020).

42. Turrigiano, G. Homeostatic Synaptic Plasticity: Local and Global Mechanisms for Stabilizing Neuronal Function. Cold Spring Harb Perspect Biol 4, a005736–a005736 (2012).

43. Wang, Y. et al. Chronic Neuronal Inactivity Utilizes the mTOR-TFEB Pathway to Drive Transcription-Dependent Autophagy for Homeostatic Up-Scaling. The Journal of Neuroscience 43, 2631–2652 (2023).

44. Wang, Y. et al. Astrocyte-secreted IL-33 mediates homeostatic synaptic plasticity in the adult hippocampus. Proceedings of the National Academy of Sciences 118, (2021).

45. Heir, R. et al. Astrocytes Are the Source of TNF Mediating Homeostatic Synaptic Plasticity. The Journal of Neuroscience 44, e2278222024 (2024).

46. Ota, Y., Zanetti, A. T. & Hallock, R. M. The Role of Astrocytes in the Regulation of Synaptic Plasticity and Memory Formation. Neural Plast 2013, 1–11 (2013).

47. Kay, R. R. Macropinocytosis: Biology and mechanisms. Cells & Development 168, 203713 (2021).

48. Pantazopoulou, M. et al. Differential intracellular trafficking of extracellular vesicles in microglia and astrocytes. Cellular and Molecular Life Sciences 80, 193 (2023).

49. Henley, J. R., Krueger, E. W. A., Oswald, B. J. & McNiven, M. A. Dynamin-mediated Internalization of Caveolae. J Cell Biol 141, 85–99 (1998).

50. Kiss, A. L. & Botos, E. Endocytosis via caveolae: alternative pathway with distinct cellular compartments to avoid lysosomal degradation? J Cell Mol Med 13, 1228–1237 (2009).

51. Pascual-Lucas, M., Fernandez-Lizarbe, S., Montesinos, J. & Guerri, C. <scp>LPS</scp> or ethanol triggers clathrin- and rafts/caveolae-dependent endocytosis of <scp>TLR</scp> 4 in cortical astrocytes. J Neurochem 129, 448–462 (2014).

52. Jiang, M. & Chen, G. Ca2+ Regulation of Dynamin-Independent Endocytosis in Cortical Astrocytes. Journal of Neuroscience 29, 8063–8074 (2009).

53. Karch, C. M. et al. A Comprehensive Resource for Induced Pluripotent Stem Cells from Patients with Primary Tauopathies. Stem Cell Reports 13, 939–955 (2019).

54. Lendemeijer, B. et al. Rapid specification of human pluripotent stem cells to functional astrocytes. Preprint at 10.1101/2022.08.25.505166 (2022).

