## Supplementary for "Neuronal autophagosomes are transported to astrocytes for degradation"

Supplementary Table 1: Overview of overnight treatments tested for LC3-transfer induction

| **Treatment** | **Mechanism** | **Concentration** |
|---|---|---|
| Rapamycin | Autophagy activation through direct mTOR inhibition | 200 nM[1] |
| BSO | Autophagy activation through increased oxidative stress | 100 µM[1] |
| KCl | Neuronal membrane depolarization triggering neuronal activation and neurotransmitter release | 50mM[2] |
| TTX | Block voltage-gated sodium channels | 1 µM[3] |
| NBQX | Competitive antagonist of AMPA receptors | 50 µM[4] |

Supplementary Table 2: Overview of endocytosis blocker

| **Endocytosis Blocker** | **Protein target** | **Pathway targeted** | **Mode of action** |
|---|---|---|---|
| Dynasore[5] | Dynamin-2 | CME/FEME, cholesterol dependent endocytosis | Prevents vesicles budding from cell membrane and reduces labile cholesterol within plasma membrane |
| PitStop2[6] | Clathrin | CME | Prevents formation of Clathrin coated pits |
| 7-Ketocholesterol[7] | PICK1 | CLIC/GEEC | interferes with actin dynamics, specifically the Arp2/3 complex |
| Cytochalasin D[8] | F-actin | Macropinocytosis/ Phagocatosis | inhibits F-actin polymerization |

Supplementary Table 3: Lentiviral Vectors

| **Vector** | **(Addgene) Ref** |
|---|---|
| Ngn2 | pLV[TetOn]-Puro-TRE3G>mNeurog2 (#198754) |
| rtTA | pLV[Exp]-EF1A>Tet3G:IRES:Neo (#198756) |
| mCherry | pHAGE-EF1a-mCHerry-IRES-Blast-WPRE (obtained from Vectorbuilder) |
| LC3-mCherry-GFP | FUW mCherry-GFP-LC3 (#110060) |
| Rat Atg5 shRNA library | 3 shRNAs ordered from VectorBuilder |

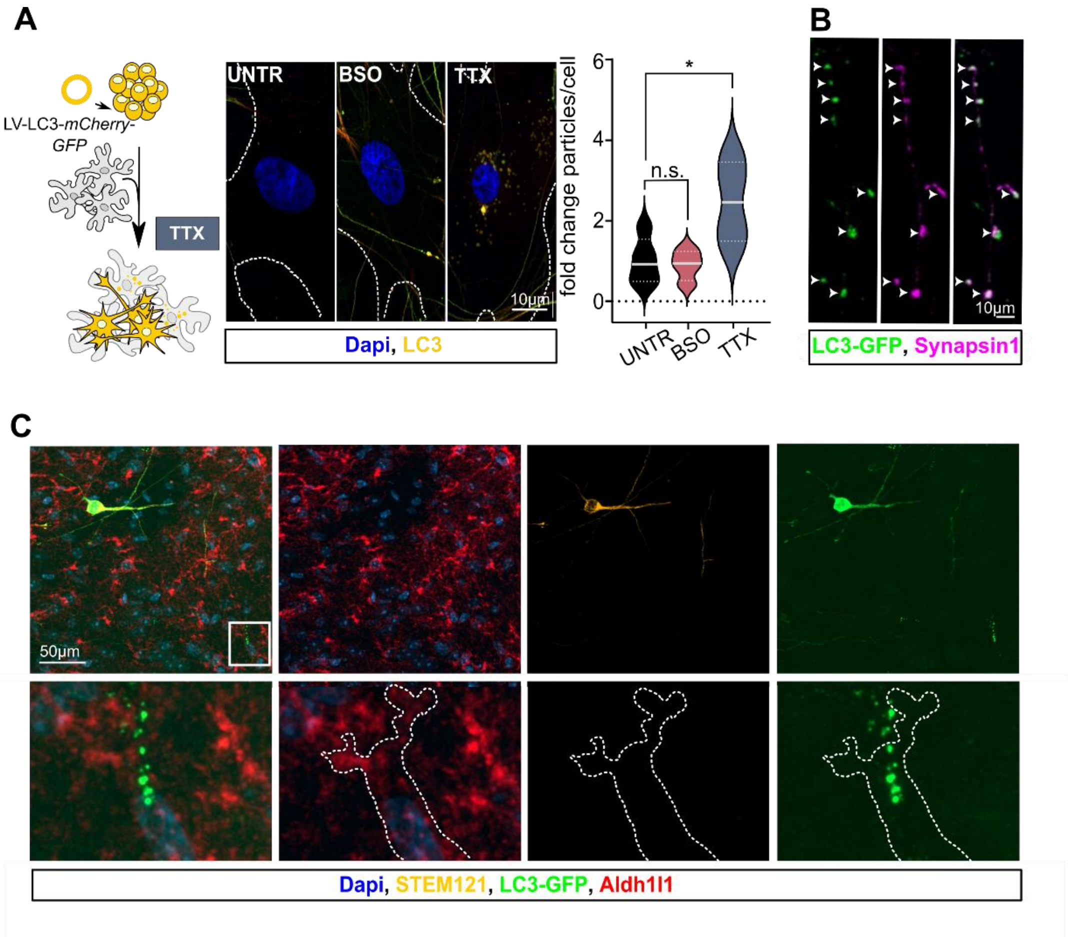


**Supplementary Figure 1: A** Schematic representation of the experimental set-up where we infected neurons with LC3-GFP-mCherry lentivirus. These iNeurons were co-cultured with unlabeled astrocytes for 4 weeks. After overnight treatment with BSO or TTX these cultures were fixed and fluorescent LC3 puncta were quantified in astrocytes. Number of puncta was compared to non-treated (UNTR) LC3-GFP-mCherry-positive cultures. N=2 **B** Representative image of a dendrite of a 4 week old iNeuron after overnight treatment with TTX. LC3-GFP puncta co-localize with the pre-synaptic marker Synapsin1 (magenta). **C** Representative image of mouse brain slice showing a STEM121-positive (yellow) neuron on the top left and LC3-GFP puncta (green) within Aldh1l1-positive (red) astrocytic structures on the bottom right.

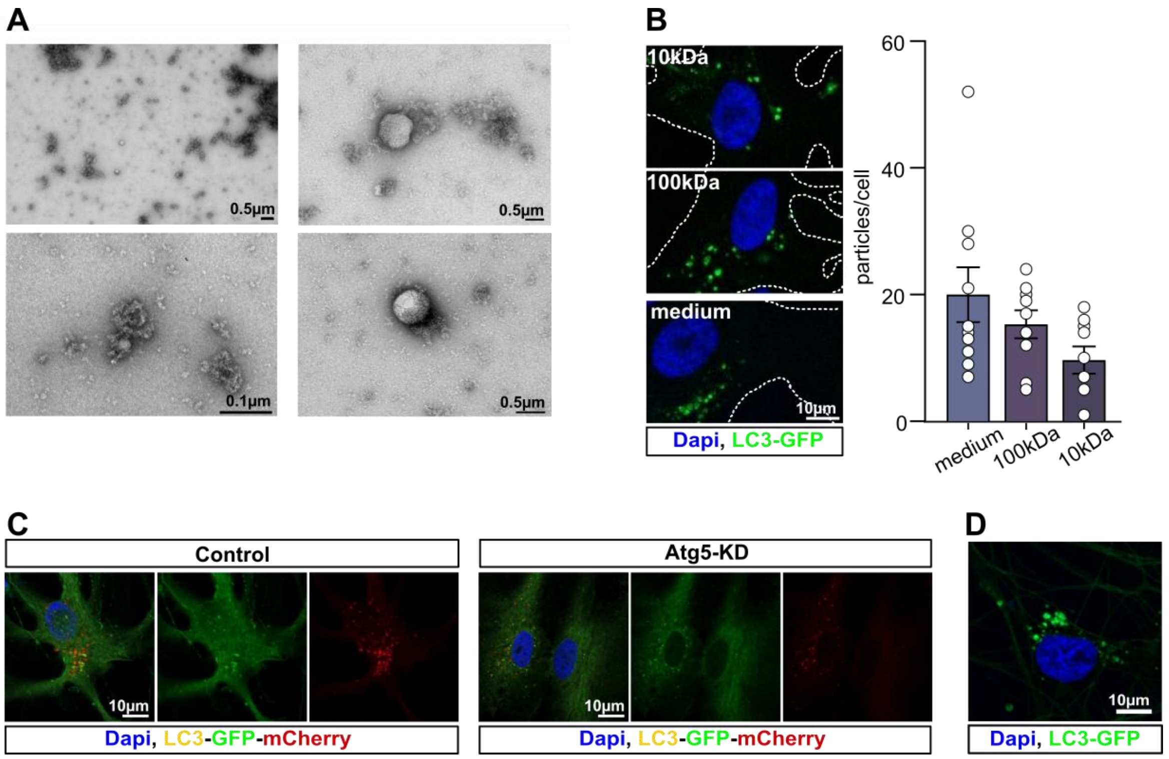


**Supplementary Figure 2: A** Representative images of negative stain EM on >100K fraction showing large protein aggregates as well as vesicles with a diameter around 500nm and smaller vesicles around 30 nm incorporated within protein aggregates. **B** Representative images of outlined astrocytes that were incubated for 3 hours with either medium of TTX treated co-cultures or the isolated >100K or >10K fractions of the same medium. LC3-GFP puncta (green) were quantified per cell. N=2 **C** Representative images of LC3-GFP-mCherry positive astrocytes that were infected with Atg5-shRNAs to block autophagosome formation in these cells. 48h after lentiviral transduction astrocytes were treated with 200 nM Rapamycin for 10 minutes. After 2 h the cells were fixed and autophagosome formation (red and green puncta) was compared to Rapamycin treated control LC3-GFP-mCherry positive astrocytes. **D** Representative image of an Atg5-KD astrocyte that was co-cultured with LC3-GFP iNeurons. 48h after shRNA transduction the culture was treated with TTX overnight and astrocytic nuclei were checked for the presence of neuronal LC3 puncta (green).

**A**

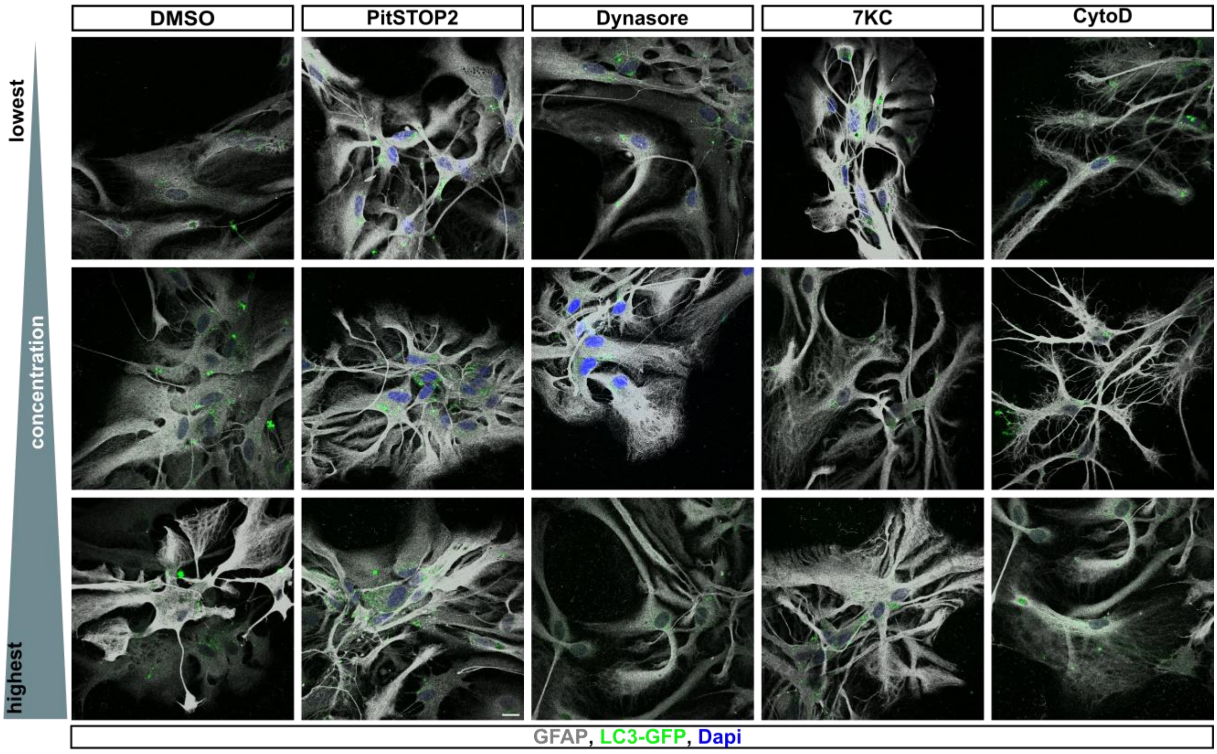


**B**

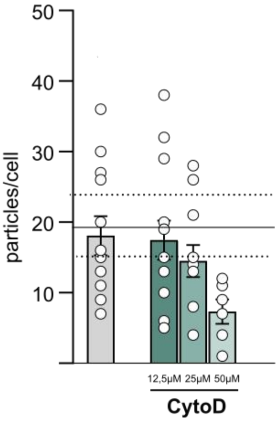


**Supplementary Figure 2: A** Representative images of Gfap (grey)- positive astrocytes that were incubated for 3 h with 12.5, 25, or 50 µM of different endocytosis blockers and medium harvested from TTX treated LC3-GFP iNeuron co-cultures. Gfap signal was used to assess effects of the different drug treatments on astrocytic health. DMSO, PitStop2, Dynasore and 7-Ketocholesterol (7KC) did not alter astrocyte structure. Cytochalasin D (CytoD), which blocks F-actin polymerization showed altered Gfap staining at the lowest applied concentration. At 25µM CytoD the Gfap staining strongly indicated that astrocyte morphology was affected. We therefore assume that CytoD treatment affected astrocytic health. **B** Quantification of LC3-GFP particles/cell in astrocytes indicated reduced LC3-GFP uptake by astrocytes incubated with 25 and 50 µM CytoD and medium of TTX treated LC3-GFP iNeuron cultures. Due to its' likely effect on general astrocytic health we decided to not draw conclusions on the effects of CytoD on macropinocytosis.

### References

1. Linda, K. *et al.* Imbalanced autophagy causes synaptic deficits in a human model for neurodevelopmental disorders. *Autophagy* 18, 423–442 (2022).

2. Füllgrabe, J. *et al.* The histone H4 lysine 16 acetyltransferase hMOF regulates the outcome of autophagy. *Nature* 500, 468–471 (2013).

3. Yuan, X. *et al.* A human in vitro neuronal model for studying homeostatic plasticity at the network level. *Stem Cell Reports* 18, 2222–2239 (2023).

4. Frega, M. *et al.* Neuronal network dysfunction in a model for Kleefstra syndrome mediated by enhanced NMDAR signaling. *Nat Commun* 10, 4928 (2019).

5. Preta, G., Cronin, J. G. & Sheldon, I. M. Dynasore - not just a dynamin inhibitor. *Cell Communication and Signaling* 13, 24 (2015).

6. von Kleist, L. *et al.* Role of the Clathrin Terminal Domain in Regulating Coated Pit Dynamics Revealed by Small Molecule Inhibition. *Cell* 146, 471–484 (2011).

7. Lyons, M. A. & Brown, A. J. 7-Ketocholesterol. *Int J Biochem Cell Biol* 31, 369–375 (1999).

8. Casella, J. F., Flanagan, M. D. & Lin, S. Cytochalasin D inhibits actin polymerization and induces depolymerization of actin filaments formed during platelet shape change. *Nature* 293, 302–305 (1981).
